# Bexarotene derivatives modify responses in acute myeloid leukemia

**DOI:** 10.1101/2021.05.17.444475

**Authors:** Gayla Hadwiger, Orsola di Martino, Margaret A. Ferris, Anh Vu, Thomas E. Frederick, Gregory R. Bowman, Peter Ruminski, Carl Wagner, John S. Welch

## Abstract

The retinoids all-trans retinoic acid (ATRA) and bexarotene are active in acute myeloid leukemia (AML), but responses beyond acute promyelocytic leukemia (APL) have been more modest than APL responses. To determine whether chemical modification of bexarotene might augment retinoid responses in AML, we screened a series of 38 bexarotene derivatives for activity in a mouse MLL-AF9 leukemia cell line, which exhibits strong synergistic sensitivity to the combination of ATRA and bexarotene. We found that RXRA potency correlated with anti-leukemic activity and that only one compound (103-4) with dual RARA/RXRA activity was capable of ATRA-independent anti-leukemic activity. We evaluated bioisostere and cyclohexane modifications for potential resistance to P450 metabolism and found that bioisosteres reduced potency and that bezopyran, cyclopentane, and cyclohexene modifications only modestly reduced susceptibility to metabolism. Collectively, these studies provide a map of the structure-activity relationships of bexarotene with outcomes related to RXRA and RARA activity, corepressor binding, compound stability, and anti-leukemic potential.

## Background

The retinoic acid receptors (RAR) and retinoid X receptors (RXR) are ligand-activated transcription factors that influence hematopoietic stem cell self-renewal and differentiation (reviewed in [1]). In normal hematopoiesis, RXRA and RARA are dynamically regulated during myeloid maturation, with highest mRNA expression in mature myeloid cells,[2] and have essential roles in granulocyte, macrophage, and osteoclast development.[3, 4, 5, 6] In acute myeloid leukemia (AML) *RARA* and *RXRA* have parallel expression among AML subtypes, with highest expression in M4/M5 myelomonocytic and monocytic subtypes.[7, 8]

Transcriptional activation or transcriptional repression by the retinoid receptors is highly dependent on the presence or absence of an activating ligand.[1, 9] RARs function as obligate heterodimers with RXRs, whereas RXRs can function either as a homodimer, or as a heterodimer with other orphan nuclear receptors (e.g. peroxisome proliferator-activated receptors (PPARs), liver X receptor (LXRs), etc). The RARA:RXRA heterodimer binds retinoic acid receptor response elements (RARE); in the absence of an activating ligand, the heterodimer is bound with a co-repressor and ligand binding induces conformational changes that displace the co-repressor and facilitate co-activator binding.[10]

ATRA and bexarotene have been explored in multiple clinical trials in non-APL forms of AML, which repeatedly suggest activity (reviewed in [11]). However, effects have generally been modest. For example, a recent study observed a median overall survival of 8.2 months in AML patients treated with ATRA + decitabine vs 5.1 months in patients receiving decitabine alone (p < 0.006), and studies using cytotoxic chemotherapy found a median overall survival of 11.3 months with the addition of ATRA vs 7.0 months without.[11-13] Likewise, bexarotene can induce maturation in AML, but clinical responses have been limited.[11, 14, 15] Subsets of patients may exhibit increased sensitivity to retinoids, and biomarker-driven trials are on-going.[11, 15]

To understand whether subtle chemical changes to bexarotene moieties might increase retinoid anti-leukemic activity achieved during RXRA activation, we evaluated a series of 38 bexarotene derivatives across assays of RXRA activation, RARA activation, anti-leukemic effects with and without concurrent ATRA, ability to release co-repressors from RARA:RXRA heterodimers, and stability of the compounds when co-cultured with HepG2 liver cells. These results provide insight into retinoid structure-activity relationships in AML cells.

## Results

### Structure and activation of RXRA

We characterized a series of 38 bexarotene derivatives for activation of RXRA in mouse MLL-AF9-derived primary leukemia cells. These cells were derived by transducing *UAS-GFP* Kit+ bone marrow cells[16] with *MSCV-MLL-AF9*. Once leukemia was established *in vivo*, these cells were harvested and transduced with *MSCV-Gal4-RXRA-IRES-mCherry*. When treated with an active RXRA ligand, mCherry+ cells become GFP+ (Schema, Figure 1A). We noted a distribution of responses, both in the EC_50_ of RXRA reporter activation and in the plateau of the median fluorescence intensity (MFI), a measure of GFP output from the reporter on an individual cell basis (Figure 1A and Supplemental Figure 1A-C). EC_50_ measured via %GFP or via GFP MFI correlated closely (R^2^ = 0.97). The MFI EC_50_ also correlated with maximum MFI, although compounds with high maximal reporter MFI tended to have low potency (i.e. high EC_50_), (R^2^ = 0.3, Figure 1B).

**Figure 1.**
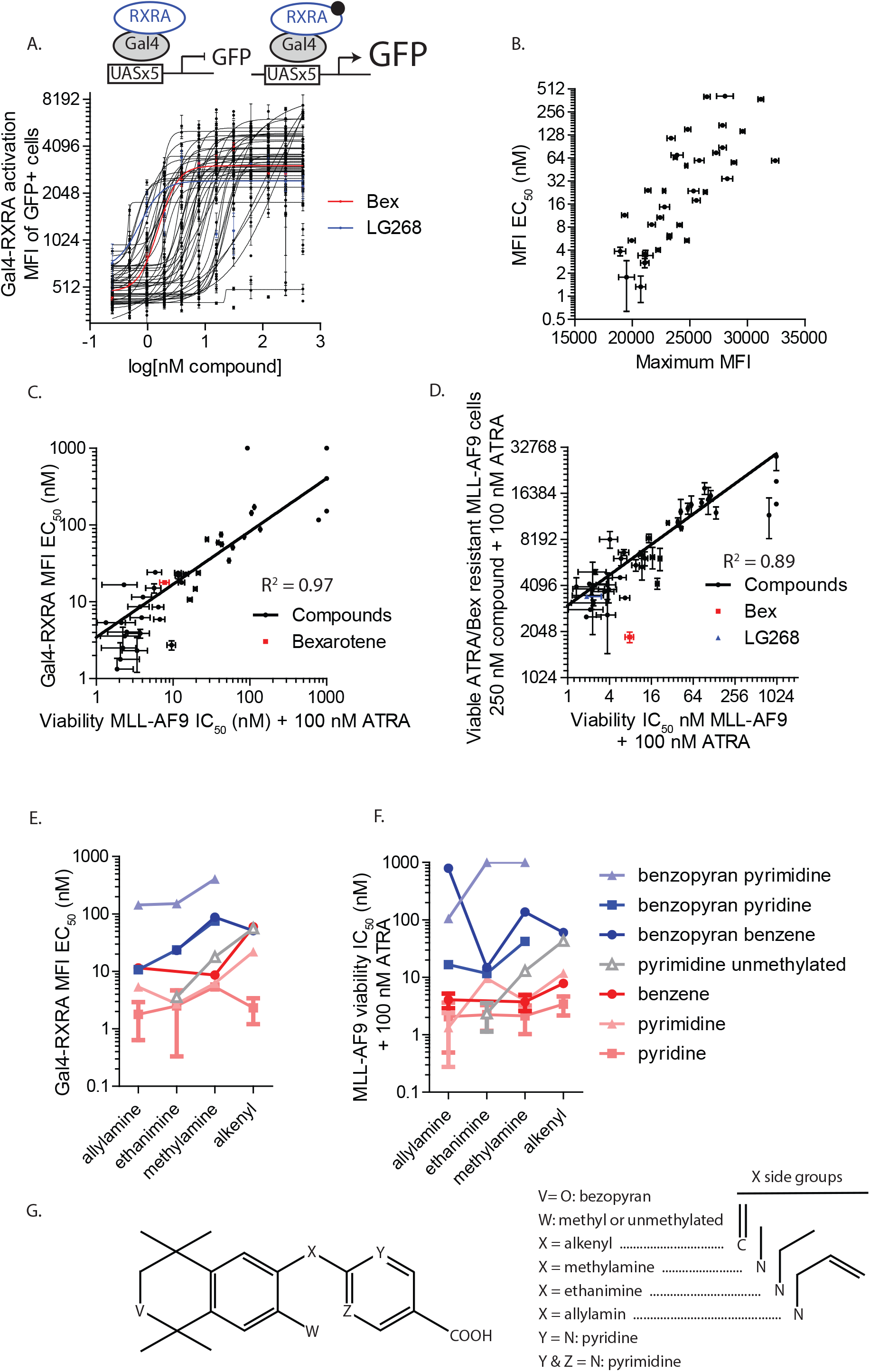
Correlation of compounds’ RXRA activity and antileukemic activity. A. Activation of a UAS-GFP/Gal4-RXRA reporter by bexarotene derivatives. B. Correlation of MFI EC_50_ and MFI plateau in UAS-GFP/Gal4-RXRA reporter output. C. Correlation of UAS-GFP/Gal4-RXRA reporter activation EC_50_ and antileukemic activity in MLL-AF9 leukemia cells. D. Correlation of antileukemic activity in parental MLL-AF9 leukemia cells vs. cells selected for retinoid resistance by culturing for 6 weeks in 100 nM bexarotene and 100 nM ATRA. E – G. Structure-activity correlations.

We had previously observed anti-leukemic synergy between RXRA ligands and RARA ligands.[8] To assess anti-leukemic effects across diverse RXRA compounds, we treated MLL-AF9 leukemia cells with increasing concentrations of each compound in the presence of 100 nM ATRA. Anti-leukemic effects were measured as the total number of viable cells after 5 days across a dose dilution to provide an IC_50_. We observed that the anti-leukemic IC_50_ correlated with the Gal4-RXRA activation EC_50_ of the compound (R^2^ = 0.97), and there was less correlation with the maximum MFI (R^2^ = 0.27) (Figure 1C). We generated an MLL-AF9 leukemia line that was resistant to ATRA and bexarotene after culturing the cells in 100 nM ATRA/bexarotene for 6 weeks. We reassessed growth after 5 days of ATRA 100 nM and 250 nM of each compound to determine whether resistance was acquired generally to the compounds or whether resistance was variable across compounds. The number of viable cells after 5 days in culture correlated with the initial IC_50_, suggesting global mechanisms of resistance rather than compound-specific sensitivity mechanisms (Figure 1D).

Within the compound set, series of molecularly related molecules could be identified for structure-activity relationships (Figure 1E-G). In general, we found that replacement of the carbon atom at C6 in the 5,6,7,8-tetrahydro-3,5,5,8,8-pentamethyl-2-naphthalenyl part of the bexarotene molecule with oxygen, to form the 3,4-Dihydro-1,1,4,4,7-pentamethyl-1H-2-benzopyran analogue, reduced both the potency and the anti-leukemic activity (i.e. they increased the GFP reporter EC_50_ and the leukemic viability IC_50_). In bexarotene, the two benzene rings are connected by a linker carbon with an alkenyl group. Substituting this alkenyl group at the linker carbon with a cyclopropyl group or replacing the carbon linker with an aliphatic substituted nitrogen as the linker (with ethyl and allyl substitution being preferred), increased both the potency and the anti-leukemic activity. Likewise, across multiple compounds, replacing the benzoic acid portion of bexarotene with nicotinic acid or a pyrimidine acid were associated with increasing potency and anti-leukemic activity.

Repeat assessments of highly potent compounds validated a series of compounds with increased potency compared with bexarotene or fluoro-bexarotene (fBex) (Supplemental Figure 1). Replicate results correlated with initial results (R^2^ = 0.83). A range of potency was noted, with the most potent compounds having similar activity as LG268. Anti-leukemic activity again correlated with RXRA activation potency (R^2^ = 0.71, Supplemental Figure 1D). Low nM EC_50_ was validated across this compound set using a second reporter assay (293T cells transfected with ApoA-Luciferase and RXRA, Supplemental Figure 1E).

### Structure and activation of RARA

We assessed all compounds for the ability to cross-activate a UAS-GFP/Gal4-RARA reporter in MLL-AF9 leukemia cells (Schema, Figure 2A). We noted several compounds with low potency RARA activation and one compound (103-4) with potent dual RXRA and RARA activity (Figure 2A, 103-4 EC_50_ Gal4-RXRA: 14 nM; Gal4-RARA: 15 nM). We determined the anti-leukemic activity of each compound in the presence and absence of concurrent ATRA, noting that 103-4 uniquely was capable of inhibiting leukemic growth independent of concurrent ATRA treatment (Figure 2B).

**Figure 2.**
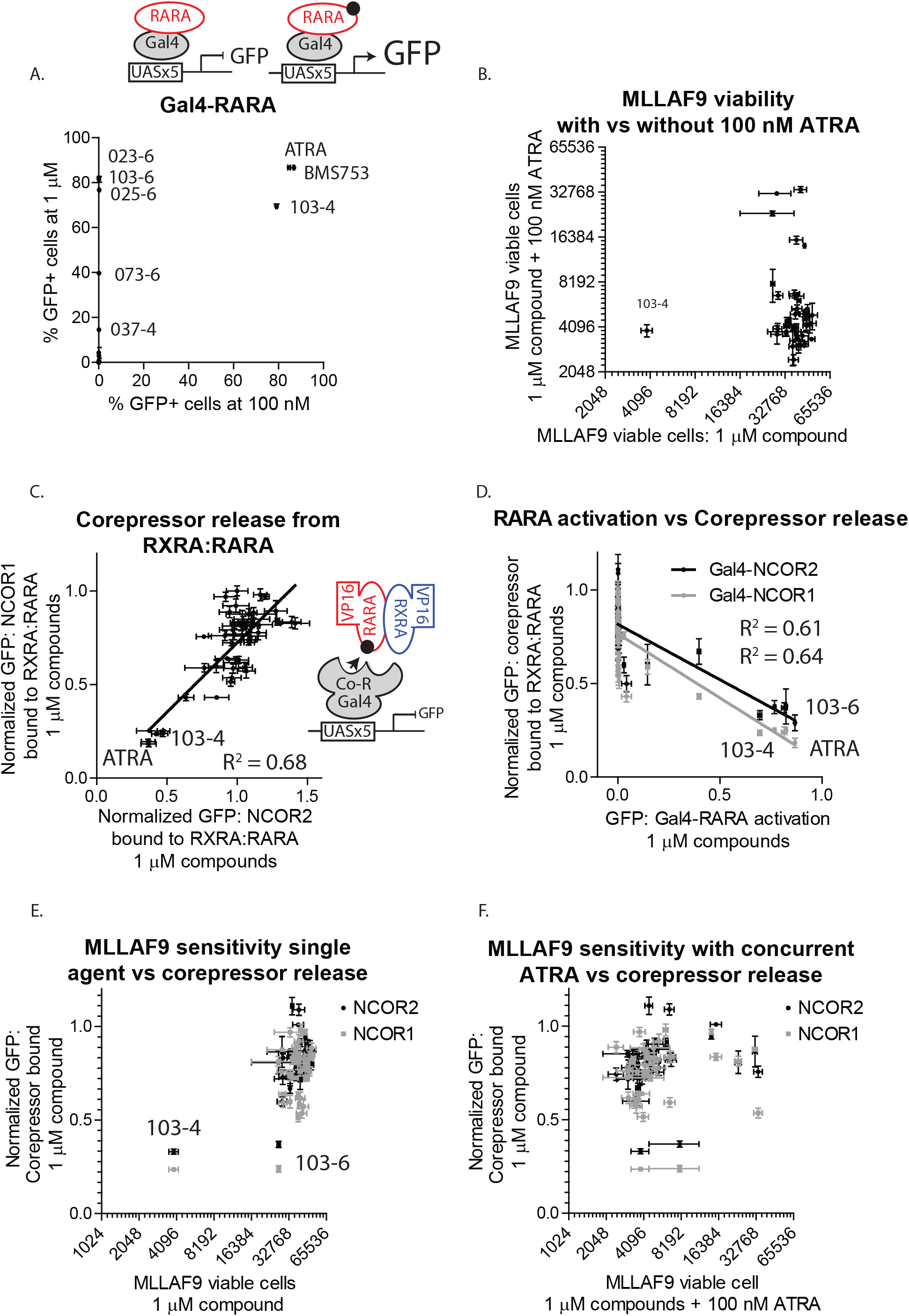
Correlation of compound RARA activity and antileukemic activity. A. Activation of UAS-GFP/Gal4-RARA reporter. B. Anti-leukemic activity of compounds with and without 100 nM ATRA. C. Mammalian two-hybrid analysis of Gal4-NCOR1 and Gal4-NCOR2 interactions with VP16-RARA:VP16-RXRA. 293T cells were transfected with UAS-GFP, VP16-RARA, VP16-RXRA, and either Gal4-NCOR1 or Gal4-NCOR2 and GFP assessed with and without compounds. D. Correlation of UAS-GFP/Gal4-RARA activation with corepressor release in mammalian two-hybrid. E – F. Correlation of corepressor release in mammalian two-hybrid assay with antileukemic activity.

RARA:RXRA forms a heterodimer, and the components of the dyad differentially interact with the corepressors NCOR1 (Nuclear Receptor Corepressor 1, formerly NCOR) and NCOR2 (Nuclear Receptor Corepressor 2, formerly SMRT).[17] We used a mammalian two-hybrid assay to determine whether different RXRA ligands differentially released co-repressors bound to RARA:RXRA heterodimers. Compound effects with Gal4-NCOR1 and RARA:RXRA correlated with effects with Gal4-NCOR2 (Figure 2C, R^2^ = 0.68). ATRA and 103-4 induced the greates co-repressor release from RARA:RXRA (Figure 2C), and these results correlated with activation of the Gal4-RARA reporter assay (Figure 2D: Gal4-NCOR1 R^2^ = 0.64; Gal4-NCOR2 R^2^ = 0.61). Thus, 103-4 uniquely exhibited both co-repressor released from RARA:RXRA and anti-leukemic activity in the absence of ATRA (Figure 2E-F).

We used *in silico* docking algorithms to assess the predicted binding of the compounds to RARA crystal structures generated in the presence of an agonist (AM580: PDB 3KMR) or an antagonist (BMS493: 3KMZ). We observed no correlation between predicted docking scores and RARA activation or co-repressor release (Supplemental Figure 2). The docking scores for the average and maximum compound conformation correlated modestly between 3KMR and 3KMZ (R^2^ _=_ 0.36 and R^2^ = 0.22, respectively).

### Prior dual RARA:RXRA ligands

Because 103-4 exhibited unique single-agent anti-leukemic activity, we evaluated the activity of two additional ligands with reported dual affinity. Compound 15b and 4-{4-[3-(trifluoromethyl)phenyl]-1,3-thiazol-2-yl}benzoic acid (PTB) had been previously suggested to maintain some dual affinity activity.[18, 19] In our assays, 15b was a potent RARA ligand, but had little RXRA activation, and was not capable of anti-leukemic activity alone or in combination with ATRA (Figure 3A-D). PTB was a weak RARA and RXRA agonist, and exhibited no anti-leukemic activity as a single agent (Figure 3E-H).

**Figure 3.**
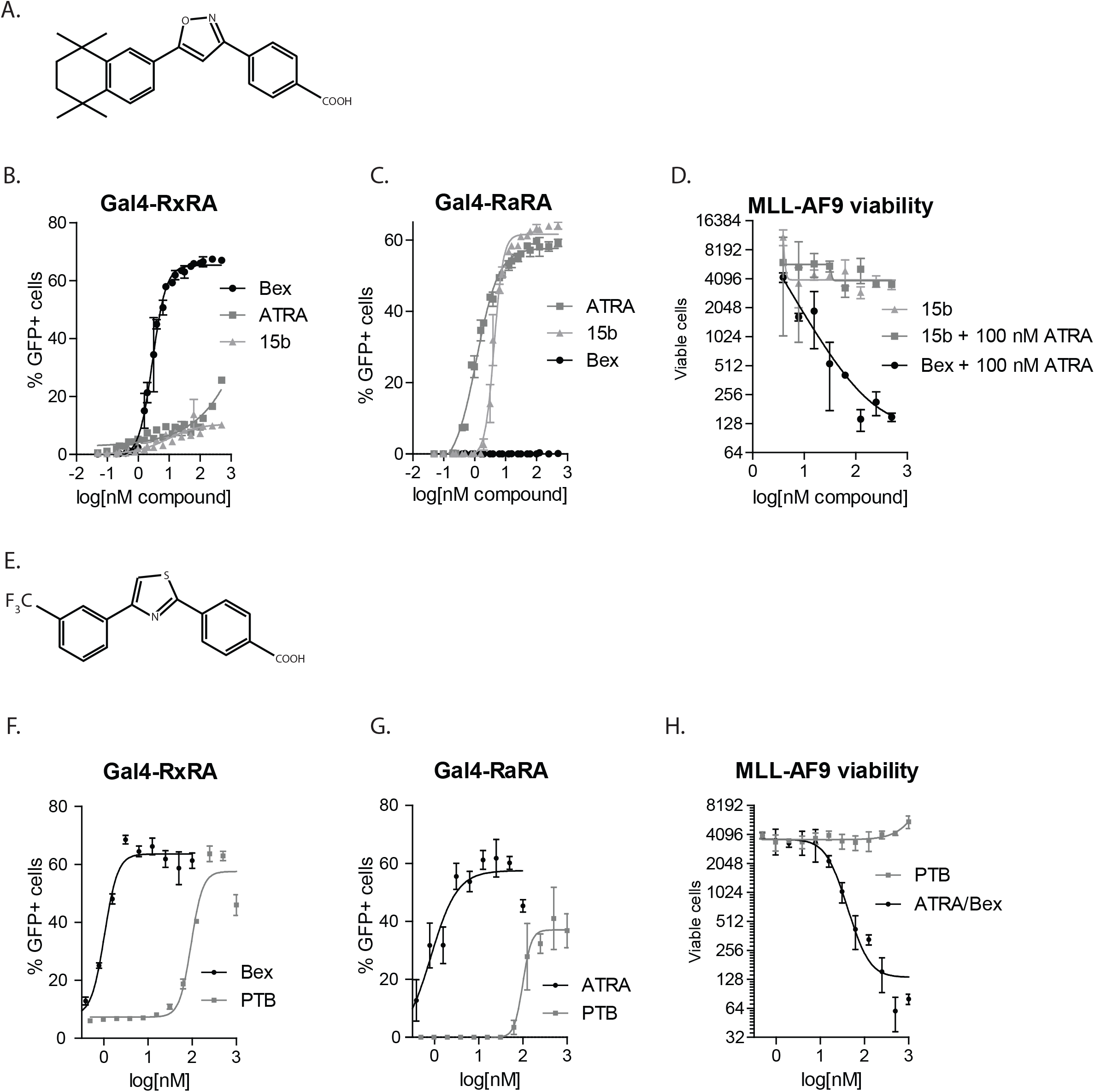
Analysis of reported dual affinity ligands. A-C. Compound 15b assessed for activation of UAS-GFP/Gal4-RXRA, UAS-GFP/Gal4-RARA, and antileukemic activity. D-F. PTB assessed for activation of UAS-GFP/Gal4-RXRA, UAS-GFP/Gal4-RARA, and antileukemic activity.

### Circumventing Metabolism

Bexarotene has a short serum half-life, in part, due to hepatic metabolism. 6- and 7-oxo and 6- and 7-hydroxy bexarotene and the corresponding hydroxylglucuronides are major metabolites in human, with glucuronidation of the carboxylic acid (acyl glucuronide) identified as metabolites as well.[20] Acid bioisosteres are simple chemical modifications of carboxylic acids that have the potential to block acyl glucuronidation while preserving the binding functionality of the free acid. Converting the benzoic acid of bexarotene to a hydroxamic acid provides the possibility of continued ionic interactions with the essential R321 that usually docks with the carboxylic acid. We synthesized a series of 3 bioisosteres (Figure 4A).[21] *In silico* docking experiments suggested variable interference with ligand binding to two different RXRA configurations (Figure 4B-C). However, all of these modifications reduced the potency RXRA activation and anti-leukemic activity of the resultant compounds (Figure 4D-E).

**Figure 4.**
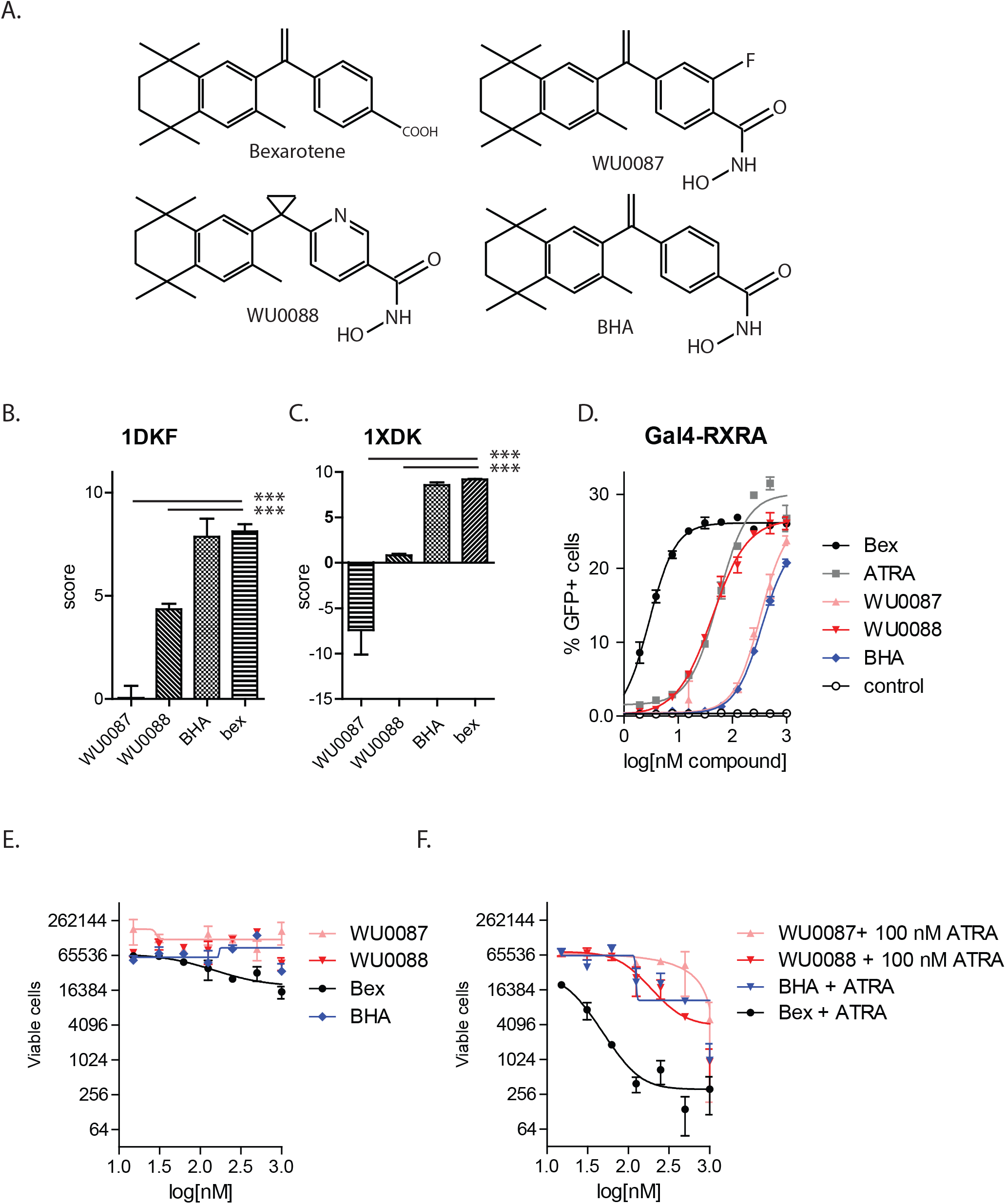
Bioisosters. A. Three bioisosteres were generated using bexarotene, fluorobexarotene, and LG268 as backbones. B-C. Docking predictions for RXRA binding using two different available crystal structures. D. Activation of UAS-GFP/RXRA-Gal4 reporter. E – F. Anti-leukemic effects without and with 100 nM ATRA.

To determine whether modifications at the C6 carbon might reduce hepatic inactivation of bexarotene, we screened the effect of HepG2 co-culture on ligand stability, comparing six such C6 modified analogs against bexarotene. Each compound was cultured for 48 or 96 hours in media alone, with HepG2 cells, or HepG2 cells that stably overexpress CYP3A4. Media was then collected and assayed for RXRA ligand activity using RXRA reporter cells (schema Figure 5A-B). The six bexarotene derivates were selected with pairwise alterations: replacing the 5,6,7,8-tetrahydro-3,5,5,8,8-pentamethyl-2-naphthalenyl part of the bexarotene molecule with the 5-membered 2,3-Dihydro-1,1,3,3-tetramethyl-1H-indene (003-6 and 007-6); replacing the C6 carbon of the 5,6,7,8-tetrahydro-3,5,5,8,8-pentamethyl-2-naphthalenyl part of the bexarotene with an oxygen to form a 3,4-Dihydro-1,1,4,4,7-pentamethyl-1H-2-benzopyran (175-6 and 169-6); or by forming an unsaturated bond between the C5 and C6 carbons of the 5,6,7,8-tetrahydro-3,5,5,8,8-pentamethyl-2-naphthalenyl part of the bexarotene molecule to form a 1,4-Dihydro-1,1,4,4-tetramethylnaphthalene (103-4 and 149-3).

**Figure 5.**
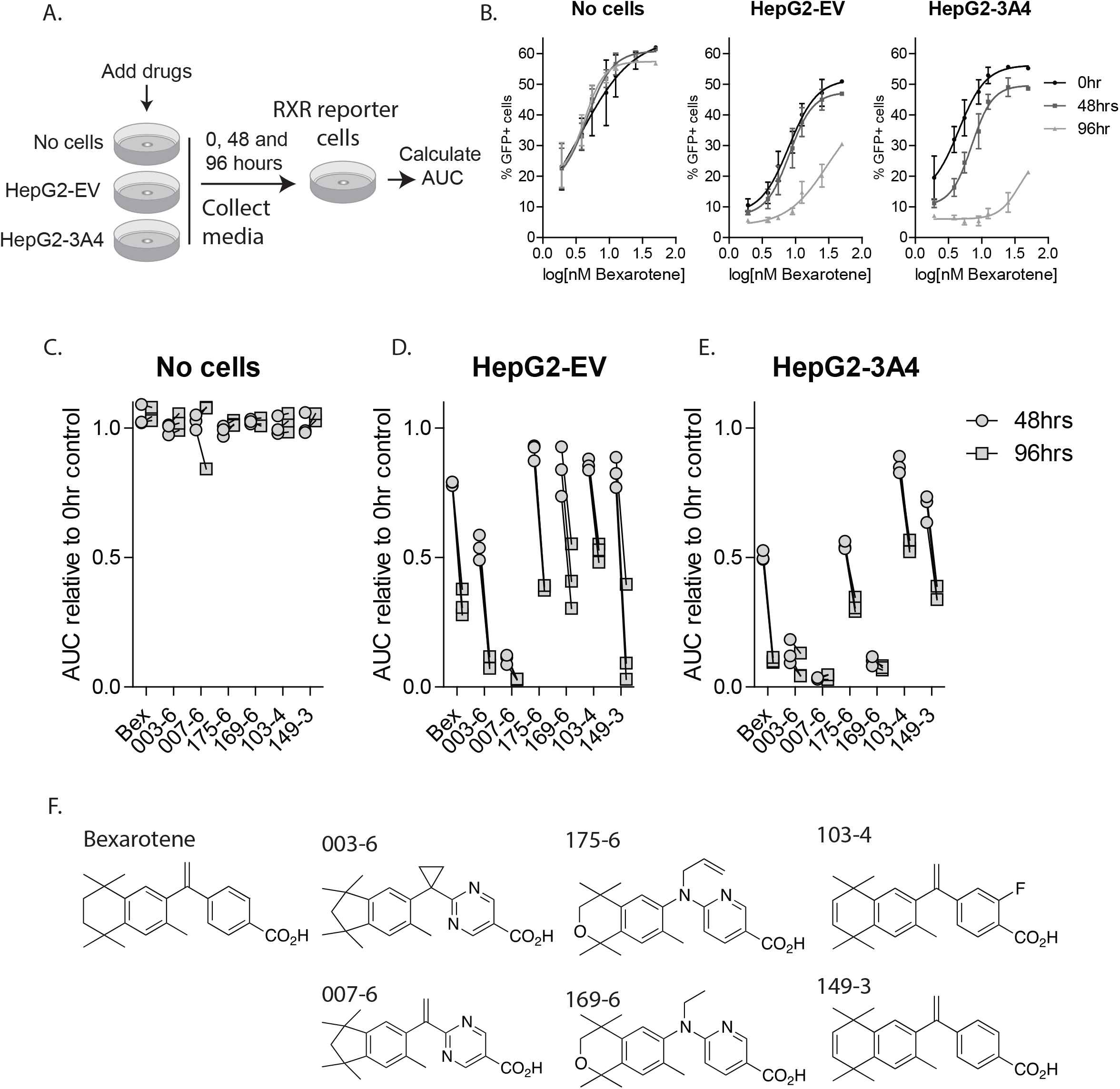
Compound metabolism by HepG2 cells. A. Schema. Compounds were added to cell culture media, with and without cells. After 0, 48, and 96 hours, media was collected and dilutions applied to UAS-GFP/Gal4-RXRA reporter cells. GFP was assessed 48 hours later and an AUC calculated. B. Representative AUC calculation. C-E. Normalized AUC when compounds were plated in media alone, with HepG2-EV cells (empty vector), or with HepG2-3A4 cells (stable expression of CYP3A4). F. Structures of compounds assessed.

All six of these compounds retained good RXR activation and anti-leukemic function (Supplemental Table 1). We noted very little reduction in RXRA activity following 96 hours of incubation in cell culture media at 37 degrees, suggesting that these compounds are remarkably stable (Figure 5C). Compared to bexarotene, the two compounds with the 5-membered 2,3-Dihydro-1,1,3,3-tetramethyl-1H-indene group (003-6 and 007-6) had increased sensitivity to HepG2A metabolism, the 3,4-Dihydro-1,1,4,4,7-pentamethyl-1H-2-benzopyran analogs had similar sensitivity (175-6 and 169-6), and the unsaturated 1,4-Dihydro-1,1,4,4-tetramethylnaphthalene analogues (103-4 and 149-3) had modestly reduced sensitivity, suggesting that they might have longer serum half-lives (Figure 5D-E).

## Discussion

The pan-RAR agonist all-trans retinoic acid (ATRA, tretinoin) transformed the treatment of APL, and has shown reproducible, but more limited activity in non-APL AML.[11-13] Bexarotene is approved for the treatment of CTCL, is also tolerable in AML patients, and has demonstrated some activity in AML, with evidence of maturation effects,[11, 14, 15] again suggesting that retinoids might be therapeutically relevant in non-APL AML.

We sought to characterize a series of bexarotene derivatives and to define their structure-activity relationships. We used a UAS/Gal4 reporter system in primary mouse leukemia cells to assess RXRA activation. The reporter reads out three parameters: percent GFP+ cells, the median fluorescence intensity of GFP among the GFP+ cells (MFI), and the plateau of the GFP MFI. When a series of dose dilutions are used, EC_50_ is calculated for the first two parameters. The %GFP+ EC_50_ and the MFI EC_50_ correlated closely (Supplemental Figure 1C). The plateau of the GFP MFI also directly correlated with the %GFP EC_50_ (Figure 1B), suggesting that less potent compounds could achieve greater GFP reporter output. Anti-leukemic IC_50_ correlated with RXRA potency (low EC_50_), not with the plateau of the GFP MFI, perhaps because less potent compounds are less toxic, and the reporter is able to achieve greater per-cell-output when exposed to compounds with low potency (i.e. high EC_50_).

Using this approach, we correlated structure and activity, noting that improved potency and anti-leukemic activity correlated with longer linker side chains, pyridine rings, and inclusion of the benzene methyl group (Figure 1).

Across this set of compounds, we noted several compounds with weak RARA activity and one compound with strong dual affinity for RARA and RXRA (Figure 2A, 103-4). Activation of RARA correlated with the release of the co-repressors NCOR1 and NCOR2 from RARA:RXRA heterodimers in a mammalian two-hybrid assay (Figure 2C-D). ATRA-independent anti-leukemic activity was exhibited by only one compound (103-4). This compound also exhibited RARA potency and the ability to release co-repressors from RARA:RXRA heterodimers, suggesting the potential necessity of RARA activation and/or co-repressor release to achieve maximum retinoid-induced anti-leukemic activity. Thus, co-repressor release from the RARA:RXRA heterodimer appears to be determined more by RARA activation than RXRA activation, consistent with prior data suggesting a subordinate role of RXRA in the RXRA:RARA heterodimer.[22, 23]

Bexarotene has a short serum half-life, in part due to hepatic metabolism.[20] We generated a series of bioisosteres to potentially block glucuronidation, and we screened compounds with modifications at the cyclohexane C6 for protection from hepatic metabolism. The bioisosteres all had marked reduction in RXRA EC_50_, likely due to steric hindrance at R321, a necessary anionic interaction to effectively dock the ligand in the ligand-binding pocket.[21] WU0087 was developed using fluorobexarotene as the backbone and proved particularly non-functional, suggesting that intramolecular interactions between the fluoro and the carbonyl groups may have formed bulky steric inhibition of the necessary interactions with R321 or other amino acid groups. Use of the more potent LG268 backbone may have helped overcome some of the detrimental effects of the bioisostere, relative to the bexarotene backbone, but this was not sufficient to generate a potent, active ligand (Figure 4).

We compared stability in tissue culture of 6 compounds against bexarotene and found that they exhibited remarkable stability in tissue culture media at 37° over 96 hours (Figure 5C). The substitution of a cyclohexene for cyclohexane modestly reduced HepG2 metabolism, whereas substitution of a cyclopentane or oxocyclohexane ring resulted in similar or more rapid metabolism by HepG2 cells (Figure 5) as well as reduced activity (Figure 1). However, these limited improvements in resistance to HepG2 metabolism are not likely to have large effects on the *in vivo* pharmacokinetics of bexarotene.

Collectively, these studies provide structure-activity relationships for bexarotene and a series of derivatives. Correlations across this broad set of chemicals demonstrate a strong relationship between potency in receptor activation assays and anti-leukemic activity, and that modifications to bexarotene can improve both the RXRA potency and the retinoid anti-leukemic activity.

## Materials and Methods

### Reagents

Bexarotene was from LC Laboratories. ATRA and PTB were from Sigma-Aldrich. Bioisostere and 15b compounds were synthesized by WuXi. Cytokines were purchased from R&D Systems. The Gal4-NCOR2, Gal4-NCOR1, NCOR2-VP16, NCOR1-VP16, VP16-RARA, and VP16-RXRA plasmids were gifts from Mitch Lazar, University of Pennsylvania. The pBABE-RXRA and ApoA1-Luciferase plasmids were gifts from Vivek Arora, Washington University. HepG2-EV and HepG2-3A4 cells were provided by the Division of Biochemical Toxicology, National Center for Toxicological Research.

### Compound synthesis

All other compounds were synthesized in the Carl Wagner laboratory.

### Generation of MLL-AF9 mouse leukemia

Mouse bone marrow Kit+ cells were isolated from *UAS-GFP* mice using an Automacs Pro (Miltenyl Biotec, San Diego, CA) per the manufacture’s protocol. Kit+ cells were plated in progenitor expansion medium (RPMI1640 medium, 15% FBS, Scf (50 ng/ml), IL3 (10 ng/ml), Flt3 (25 ng/ml), Tpo (10 ng/ml), L-glutamine (2 mM), sodium pyruvate (1 mM), HEPES buffer (10 mM), penicillin/streptomycin (100 units/ml), β-mercaptoethanol (50 µM)) overnight and transduced by spinfection with 10 µg/ml polybrene and 10 mM HEPES at 2400 rpm, 30°C for 90 minutes in an Eppendorf 5810R centrifuge. Cells were transplanted into sublethally irradiated mice and subsequent leukemia harvested 2-4 months later, as expected.[24] *MLL-AF9* leukemia cells were cultured *in vitro* using similar media, but without Flt3, or Tpo. Fluorescence was detected on an Attune NxT Flowcytometer (Invitrogen) or ZE5 Cell Analyzer (Biorad).

### Mice

*UAS-GFP* mice were bred as described.[6, 16] The Washington University Animal Studies Committee approved all animal experiments.

### Retrovirus production

Retrovirus production was performed as described using previously described vectors.[6, 16] 7 × 10^6^ 293T/17 cells were seeded in a 150 cm^2^ dish in DMEM (high glucose) + 10% FBS + 1% Glutamax, 18-24 hours before transfection and grown to 80% confluence. 30 µg of DNA, 21.5 µg Ecopak, and 1.25 mL of DMEM were mixed. 40 µL Lipofectamine 3000 + 1.25 mL DMEM were mixed. The two mixtures were incubated for 5 minutes, then mixed together and incubated for 15 minutes. 1.25 mL of the mixture were dropped-wise onto the 293T/17 cells. Fresh medium was changed after 18-24 hours transfection. Virus was collected at 48 hours and 72 hours and concentrated with Lenti-X Concentrator (Clontech). The Virus was resuspended in DMEM and stored at -80C.

### EC_50_ and IC_50_ calculations

UAS-GFP/MLL-AF9 leukemia cells were transduced with either MSCV-3xFlag-Gal4 (DBD)-RXRA (LBD)-IRES-mCherry or MSCV-Gal4 (DBD)-RARA (LBD)-IRES-mCherry. Typically, compounds were diluted in 100 µls in 96 well plates in duplicate, starting at 1 µM through 12 1:1 dilutions. Cells were added in 100 µl media aliquots and then evaluated by flow cytometry 48 hours later. Cell viability was similarly plated on day 1. On day 3, 10 µl were replated into freshly diluted compounds and new media. On day 5 total viable cells per well were evaluated by flow cytometry. EC_50_ and IC_50_ were calculated using Prism Graphpad.

### Mammalian two-hybrid assay

293Tcells were co-transfected using Lipofectamine 3000 (Invitrogen) with plasmids encoding the reporter: *UAS-GFP; Gal4-*fusion vectors as “bait”; and *VP16-*fusions as “prey”. The percentage of GFP+ cells was assessed 48 hours after transfection by flow cytometry.

### Luciferase detection

293T cells were transfected with pBABE-RXRA in combination with ApoA1-Luciferase using Lipofectamine 2000 (Invitrogen). Six hours after transfection, the cells were collected and plated into a 48 well plate in 1% BSA media and treated in triplicate. After 40 hours incubation, the cells were harvested and assayed for luciferase (Luc Assay System with Reporter Lysis Buffer, Promega) in a Beckman Coulter LD400 plate reader.

### HepG2 analysis

HepG2 cells that stably express cytochrome P450s (CYPs) and a control, empty vector (EV), with a Blasticidin marker were grown in the recommended media (HepG2 Media: Gibco High Glucose DMEM with 10% FBS, 100U/mL Pen/Strep, 0.25 ug/mL Blasticidin, 1% Glutamax, 1 mL NEAA (non-essential amino acids)). The cells were plated onto a 12 well plate @ 500,000 cells per well in 0.5 mL media and grown overnight at 37°C in a tissue culture CO2 incubator. The next day, the media was replaced with new media, without Pen/Strep, and drug dilutions were added. Media for each drug dilution was collected from the cells at 0, 48, and 96 hours and frozen @ -80°C. UAS-GFP MLL-AF9 reporter cells were plated onto a 96 well plate at 10,000 cells per well in 0.1 mL media from the thawed HepG2 metabolized drug aliquots via serial dilutions. After 48 hours, the cells were analyzed on an Invitrogen Attune NxT flow cytometer.

### Compound docking

Analysis was performed with Surflex-dock (26). MOL2 files for each compound were generated from SMILES strings using Open Babel.(41) Surflex-Dock receptor protomols were generated with a threshold of 0.25 and a bloat of 2.0 and subsequently docked using the default ‘-pgeom’ docking accuracy parameter set.

### Data analysis

Flow cytometry data was analyzed with FlowJo software version 10. Statistical analysis was performed using Prism (Graphpad). T-test and ANOVA tests were performed, as appropriate. Error bars represent standard deviation. Data points without error bars have standard deviations below Graphpad’s limit to display.

## Acknowledgments

We thank the Alvin J. Siteman Cancer Center at Washington University School of Medicine and Barnes-Jewish Hospital in St. Louis, MO. for the use of the Flow Cytometry Core. The Siteman Cancer Center is supported in part by an NCI Cancer Center Support Grant P30 CA91842. We thank Conner York for technical assistance. This work was supported by NIH R01 HL128447 (JS Welch), by the Siteman Investment Program (JS Welch), the Washington University SPORE DRP (JS Welch), and the Children’s Discovery Institute (JS Welch). Author Contributions: J.S.W. and G.H. designed experiments, performed experiments, and wrote the manuscript. A.V., O.d.H, M.A.F., T.E.F, G.R.B, P.R. designed and performed experiments. Compounds were synthesized by C.W.

---

**Supplemental Figure 1.** Assessment of compounds with EC_50_ potency greater than bexarotene. A. Activation of UAS-GFP/Gal4-RXRA reporter in MLL-AF9 leukemia cells measured by percent GFP+ cells. B. Activation of UAS-GFP/Gal4-RXRA reporter in MLL-AF9 leukemia cells measured by median fluorescence intensity (MFI) of GFP+ cells. C. Correlation of EC_50_ calculated by percent GFP+ and by MFI. D. Correlation of antileukemic activity, measured by IC_50_ of each compound, and the EC_50_ activation of the UAS-GFP/Gal4-RXRA reporter. E. Activation of a ApoA-luciferace reporter. F. Structures of labeled, potent compounds.

**Supplemental Figure 2.** Predicted compound binding to RARA. A. Docking scores for compounds bound to 3KMR (RARA bound to the RARA agonist AM580). B. Docking scores for compounds bound to 3KMR (RARA bound to the RARA antagonist BMS493). Each dot represents results for a separate compound conformation. Red indicates compounds with activity in UAS-GFP/Gal4-RARA reporter assay (Figure 2A).

