## Supplementary figures and images for "Bexarotene derivatives modify responses in acute myeloid leukemia"

### Supplemental Figure 1

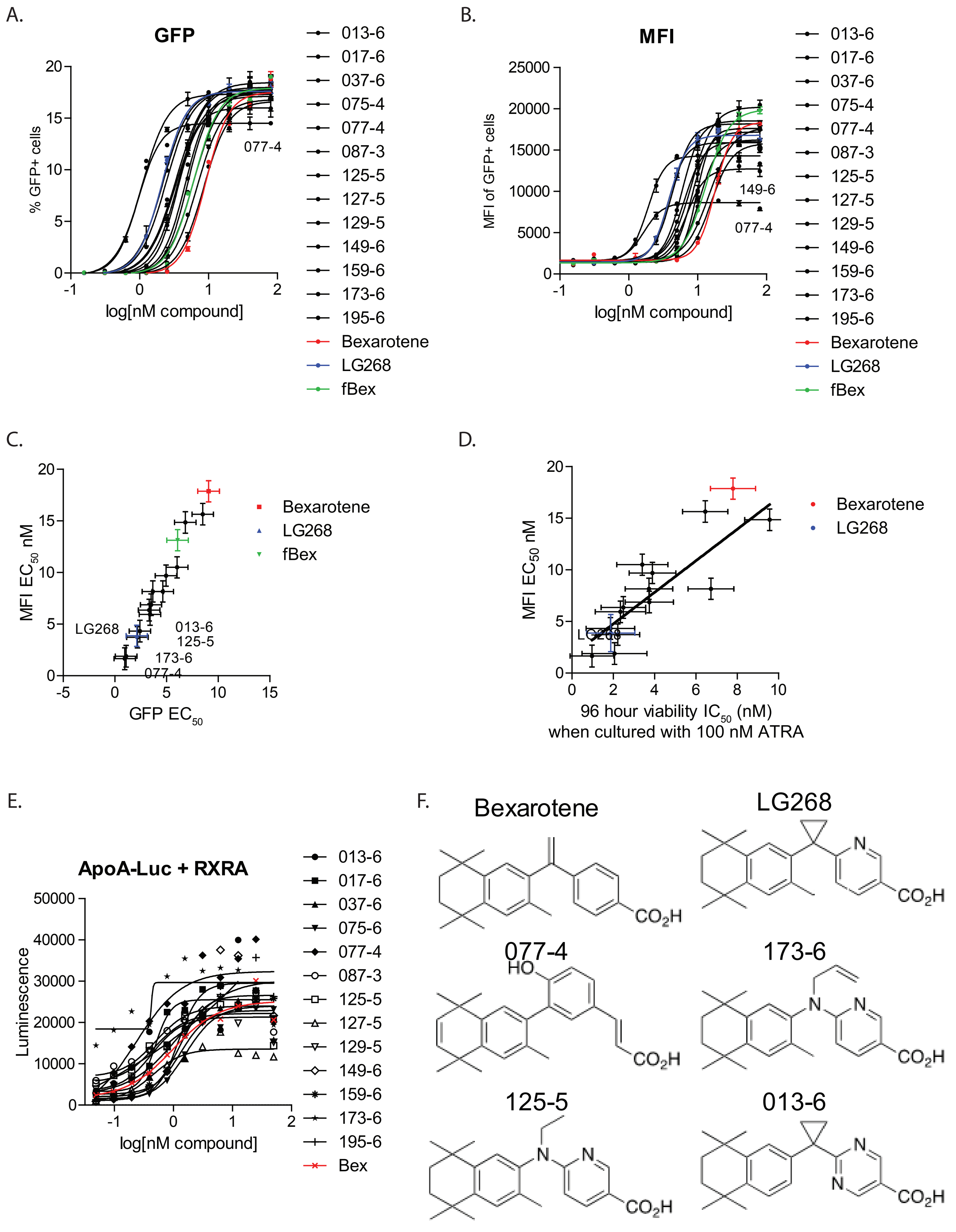


Supplemental Figure 1.
