## Supplemental Figure 2 for "Bexarotene derivatives modify responses in acute myeloid leukemia"

A.

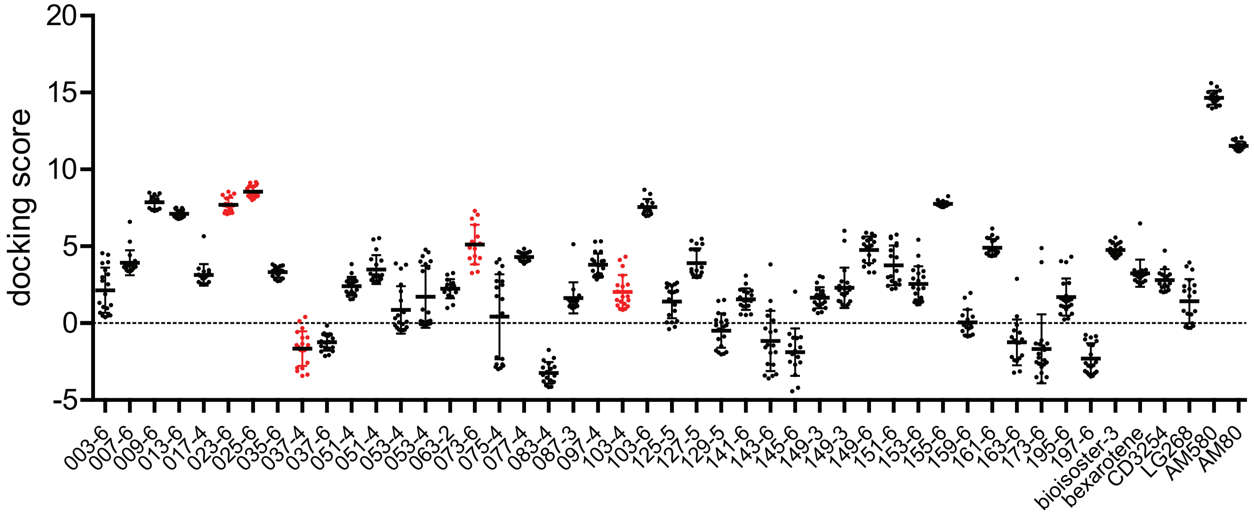


**Docking to crystal structure 3KMR - RARA bound with AM580 (agonist)**

B.

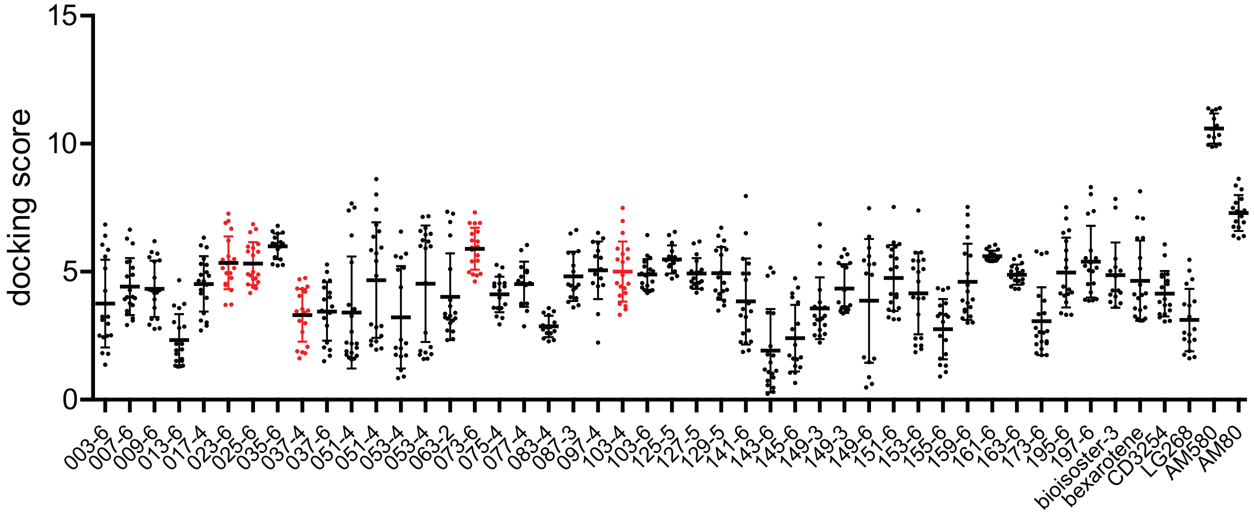


**Docking to crystal structure 3KMZ - RARA bound with BMS493 (antagonist)**

Supplemental Figure 2.
